# Structural basis of RfaH-mediated transcription-translation coupling

**DOI:** 10.1101/2023.11.05.565726

**Authors:** Vadim Molodtsov, Chengyuan Wang, Jason T. Kaelber, Gregor Blaha, Richard H. Ebright

## Abstract

The NusG paralog RfaH mediates bacterial transcription-translation coupling on genes that contain a DNA sequence element, termed an *ops* site, required for pausing RNA polymerase (RNAP) and for loading RfaH onto the paused RNAP. Here we report cryo-EM structures of transcription-translation complexes (TTCs) containing RfaH. The results show that RfaH bridges RNAP and the ribosome, with the RfaH N-terminal domain interacting with RNAP, and with the RfaH C-terminal domain interacting with the ribosome. The results show that the distribution of translational and orientational positions of RNAP relative to the ribosome in RfaH-coupled TTCs is more restricted than in NusG-coupled TTCs, due to the more restricted flexibility of the RfaH interdomain linker. The results further show that the structural organization of RfaH-coupled TTCs in the "loading state," in which RNAP and RfaH are located at the *ops* site during formation of the TTC, is the same as the structural organization of RfaH-coupled TTCs in the "loaded state," in which RNAP and RfaH are located at positions downstream of the *ops* site during function of the TTC. The results define the structural organization of RfaH-containing TTCs and set the stage for analysis of functions of RfaH during translation initiation and transcription-translation coupling.

**One sentence summary:** Cryo-EM reveals the structural basis of transcription-translation coupling by RfaH.

## Introduction

Bacterial transcription and bacterial translation occur in the same cellular compartment and occur at the same time^1-2^. In most bacteria, including the standard model organism *Escherichia coli*, transcription and translation are functionally coordinated processes, in which the rate of transcription by the first molecule of RNA polymerase (RNAP) synthesizing an mRNA is coordinated with the rate of translation by the first ribosome ("lead ribosome") translating the mRNA^2-13^. Transcription-translation coupling is thought to have important mechanistic consequences for transcription, translation, and transcriptional and translational regulation^3-23,^. The physical coupling of an RNAP molecule and a lead ribosome is hypothesized to increase transcription elongation rates and to suppress transcription pausing, by enabling a slowly transcribing or paused RNAP molecule to be "pushed" by a lead ribosome^3-17^, and is hypothesized to suppress transcription termination by sterically blocking the formation of termination RNA hairpins and sterically blocking the function of termination factor Rho^3-13,18-23^. The physical coupling of an RNAP molecule and a lead ribosome also plays roles in classic mechanisms for regulation of bacterial gene expression, including transcription attenuation^12,16-17^ and riboswitch function^11-12^.

Coordination between transcription and translation in *E.coli* is mediated by transcription elongation factors of the NusG/RfaH family^4-13,24-28^. NusG serves as a general coupling factor that couples transcription and translation at many genes in *E.coli*^4-13,24-28^. NusG paralog RfaH serves as a specialized regulon-specific coupling factor that couples transcription and translation at a subset of genes in *E. coli* that contain a specific DNA sequence element, termed an *ops* site, required for pausing RNAP and for loading RfaH onto the paused RNAP^5-7,10-12,24-28,^. NusG and RfaH each consist of an N-terminal domain (NusG-N and RfaH-N) that interacts with RNAP, a C-terminal domain (NusG-C and RfaH-C) that interacts with ribosomal protein S10, and an interdomain linker^4-5,7-13,24-28^. Transcription-translation coupling in *E. coli* is further modulated by a second transcription elongation factor: NusA^29^.

Structures of NusG-containing *E.coli* transcription-translation complexes (TTCs) recently have been reported^30-32^. The results define two distinct classes of TTCs: (i) "collided TTCs", which cannot accommodate NusA, and which have structural features that suggest they are translationally inactive, or only partly active, products of collision between a ribosome and RNAP (TTC-A: Fig. 1a, left) ^30-32^, and (ii) "coupled TTCs", which can accommodate NusA, and which have structural features that suggest they are translationally active complexes that mediate functional transcription-translation coupling (TTC-B; Fig. 1a, center and right) ^31-32^ (Fig. 1c).

**Fig. 1.**
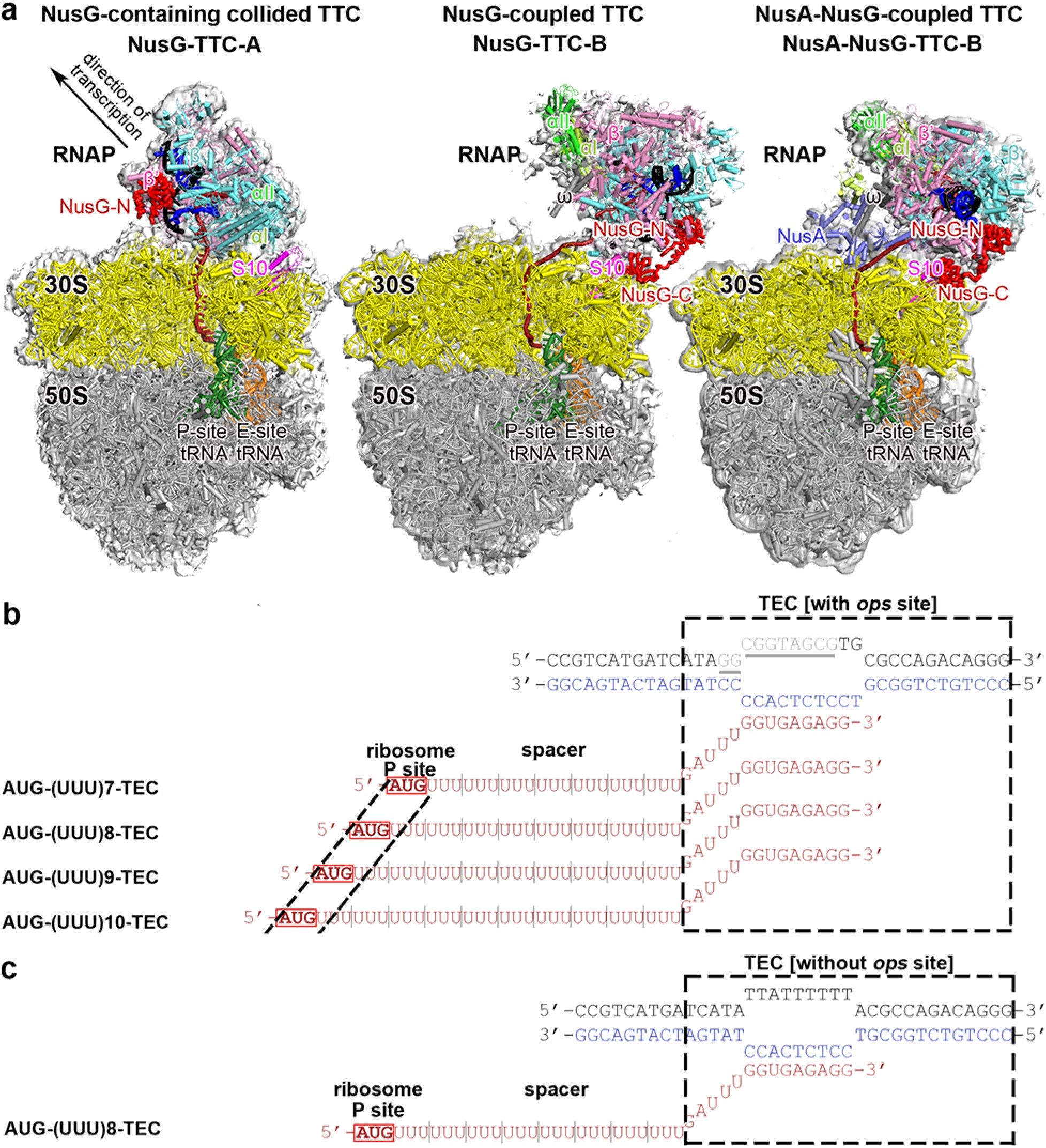
Structures of NusG-containing TTCs and nucleic-acid scaffolds for structure determination of RfaH-containing TTCs. **(a)** NusG-containing TTCs. Left: In NusG-containing collided TTC, NusG makes no interactions with ribosome (NusG-TTC-A; PDB )^31-32^. Center: In NusG-coupled TTC, NusG bridges TEC and ribosome (NusG-TTC-B; PDB 6XII)^31-32^. Right: In NusA-, NusG-coupled TTC, NusG bridges TEC and ribosome, and NusA forms second bridge between TEC and ribosome. (NusA-NusG-TTC-B; PDB 6X6T)^32^. Images show EM density (gray surface) and fit (ribbons) for TEC, NusG, and NusA (at top; direction of transcription indicated by arrow in the left panel and directly toward viewer in center and right panels) and for ribosome 30S and 50S subunits and P- and E-site tRNAs (at bottom). RNAP β’, β, α^I^, α^II^, and ω subunits are in pink, cyan, light green, dark green, and gray; 30S subunit, 50S subunit, P-site tRNA, and E-site tRNA are in yellow, gray, green, and orange; DNA nontemplate strand, DNA template strand, and mRNA are in black, blue, and brick-red. NusG, NusA, and ribosomal protein S10 are in red, light blue, and magenta. Rbosome L7/L12 stalk omitted for clarity in this and subsequent images. **(b)** Nucleic-acid scaffolds for structure determination of RfaH-containing collided TTC (RfaH-TTC-A) and RfaH-coupled TTCs in the loading state (RfaH-TTC-B^loading^). Each scaffold comprises nontemplate-strand oligodeoxyribonucleotide (black, with *ops*-site nucleotides in gray and underlined), template-strand oligodeoxyribonucleotide (blue), and one of four oligoribonucleotides having spacer lengths, n, of 7, 8, 9, and 10 codons, corresponding to mRNA (brick red). Dashed black box labeled "TEC," portion of nucleic-acid scaffold that forms TEC upon addition of RNAP (10 nt nontemplate- and template-strand ssDNA segments forming "transcription bubble," 10 nt of mRNA engaged with template-strand DNA as RNA-DNA "hybrid," and 5 nt of mRNA, on diagonal, in RNAP RNA-exit channel); dashed black lines labeled "ribosome P-site," mRNA AUG codon intended to occupy ribosome active-center P site upon addition of ribosome and tRNA^fMet^; "spacer," mRNA spacer between TEC and AUG codon in ribosome active-center P site. **(c)** Nucleic-acid scaffolds for structure determination of RfaH-coupled TTCs in loaded state (RfaH-TTC-B^loaded^). Colors as in b.

In NusG-containing coupled TTCs, NusG bridges RNAP and the lead ribosome, with NusG-N interacting with RNAP, and with NusG-C simultaneously interacting with ribosomal protein S10 in the ribosome 30S subunit (Fig. 1a, center and right)^31-32^. NusA forms a second bridge between RNAP and the lead ribosome (Fig 1a, right)^32^; this second bridge supplements and supports, but cannot substitute for, the bridge formed by NusG^32^.

It has been hypothesized that specialized coupling factor RfaH, like the general coupling factor NusG, bridges RNAP and the lead ribosome, with RfaH-N interacting with RNAP, and with RfaH-C simultaneously interacting with the ribosomal protein S10 in the ribosome 30S subunit. ^5-7,10-12,24-28^ Consistent with this hypothesis, RfaH-N has been shown to interact with RNAP^5,33^, interacting with the same site on RNAP as NusG-N^5,33^, and RfaH-C has been shown to interact with ribosomal protein S10, interacting with the same part of ribosomal protein S10 as NusG-C^5^. However, three key questions about RfaH have remained unanswered. First, what is the structure of the RfaH-coupled TTC? Second, what are the differences, if any, between the structures of the RfaH-coupled TTC and the NusG-coupled TTC? Third, what are the differences, if any, between the structures of the RfaH-coupled TTC in the "loading state," in which RNAP and RfaH are located at the *ops* site within a transcription unit during formation of the TTC, and the RfaH-coupled TTC in the "loaded state," in which RNAP and RfaH are located at positions downstream of the *ops* site during subsequent function of the TTC?

Here, we report cryo-EM atomic structures of RfaH-containing collided TTCs and coupled TTCs, analyzing complexes both in the "loading state" and in the "loaded state," and analyzing complexes both in the absence of NusA and in the presence of NusA.

### Structure determination

We analyzed synthetic nucleic-acid scaffolds similar to those used previously to determine structures of NusG-containing collided TTCs and coupled TTCs (Fig. 1A-B)^32^. The scaffolds comprise three parts: (i) DNA and mRNA determinants that direct formation of a transcription elongation complex (TEC) upon interaction with RNAP, (ii) an mRNA AUG codon that enables formation of a translation complex having the AUG codon positioned in the ribosome active-center product site (P-site) upon interaction with a ribosome and tRNA^fMet^, and (iii) an mRNA spacer having a length, *n*, of 7, 8, 9, or 10 codons (21, 24, 27, or 30 nt) between (i) and (ii) (Fig. 1b-c). For determination of structures of RfaH-containing TTCs in the "loading state," the scaffolds contained an *ops* site, with a consensus sequence and consensus positioning relative to the unwound transcription bubble (Fig. 1b); for structures of RfaH-containing TTCs in the "loaded state," the *ops* site was replaced by an unrelated sequence (Fig. 1c). For each scaffold, we incubated the scaffold with RNAP to form a TEC; then incubated with RfaH, or both RfaH and NusA, to form an RfaH-containing or NusA-RfaH-containing TEC; and then incubated with ribosomes and tRNA^fMet^ to form an RfaH-containing or NusA-RfaH-containing TTC. For each resulting complex, we then determined the structure by use of single-particle reconstruction cryo-EM (Figs. S1-S5; Table S1).

### Structure of RfaH-containing collided TTC

The structure of an RfaH-containing collided TTC (RfaH-TTC-A) was obtained using an *ops*-containing scaffold having an mRNA spacer of 7 codons (Figs. 2, S1; Table S1). In the structure, the TEC is oriented relative to the ribosome such that RNA proceeds directly from the TEC RNA-exit channel into the ribosome S30 subunit mRNA-entrance channel, and, from there, proceeds to the ribosome active-center P-site (Fig. 2). In the structure, RfaH-N is present, is fully ordered, makes protein-protein interactions with the RNAP β’ and β pincer tips and the RNAP lobe that are identical to those in the structure of RfaH-TEC complexes in the absence of ribosomes^33^, and makes sequence-specific protein-DNA interactions with *ops*-site DNA in the unwound transcription bubble region that are identical to those in the structure of RfaH-TEC complexes in the absence of ribosomes^33-34^ (Figs. 2a, S1f). RfaH-C is disordered, indicating that it makes no stable specific interactions, and, instead, exists as an ensemble of different conformational states. The spatial relationship of the TEC relative to the ribosome in the RfaH-containing collided TTC is identical to that in structures of collided TTCs obtained in the absence of coupling factors^30,32^ and in structures of collided TTCs obtained in the presence of coupling factor NusG^31-32^.

**Fig. 2.**
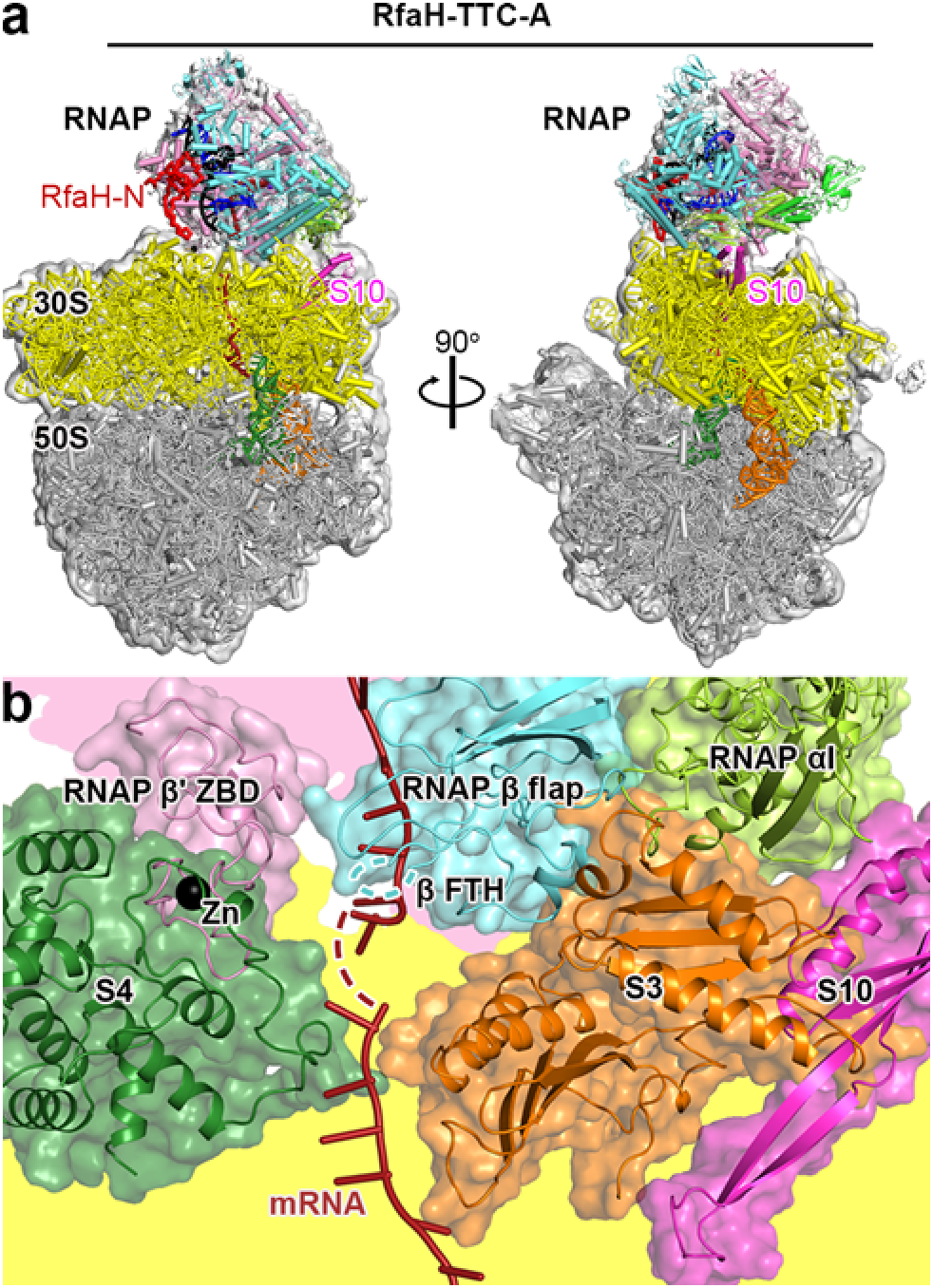
Structure of RfaH-containing collided TTC. **(a)** Structure of RfaH-TTC-A (n = 7; Table S1). Two orthogonal views. RfaH, red. Other colors as in Fig. 1a. **(b)** RNAP-ribosome interface in RfaH-TTC-A, showing RNAP β’ zinc binding domain, (ZBD, pink; Zn^2+^ ion as black sphere), RNAP β flap, cyan, RNAP β flap tip helix (β FTH; disordered residues indicated by cyan dashed line), and RNAP α^I^ (green) interacting with ribosomal proteins S4 (forest green), S3 (orange), and S10 (magenta) and with mRNA (brick red). Portions of RNAP β’ and ribosome 30S not involved in interactions are shaded pink and yellow, respectively.

The RNAP-ribosome interface in the RfaH-containing collided TTC is large, involving protein-protein interactions by the RNAP β’ zinc-binding domain (ZBD), the RNAP β flap tip (FT), and the RNAP α C-terminal domain (αCTD) with, respectively, ribosomal proteins S4, S3, and S3 in the ribosome 30S head and 30S body (Fig. 2b). This interface, like the essentially identical interfaces present in collided complexes formed in the absence of coupling factors^30,32^ or in the presence of NusG^31-32^, cannot accommodate transcription elongation factor NusA, and is expected, because it spans the ribosome 30S head and 30S body, to inhibit the ribosome 30S head swivelling and unswivelling that occurs in each translation cycle in processive translation^35-37^, and thus is expected to be inactive in, or only partly active in, processive translation. We infer that the RfaH-containing collided TTC is a complex that occurs only upon collision of a lead ribosome with RNAP and is not a complex that mediates processive coupled transcription-translation.

### Structures of RfaH-containing coupled TTCs in the loading state

Structures of RfaH-containing coupled TTCs in the loading state (RfaH-TTC-B^loading^) were obtained using the nucleic-acid scaffold having an mRNA spacer of 7 codons (as a second subclass, in addition to the subclass comprising the RfaH-containing collided TTC discussed above; Figs. 3a-d; S1; Table S1) and also were obtained using nucleic-acid scaffolds having mRNA spacers of 8, 9, or 10 codons (Figs. 3a-e; S2; Table S1). The structural organization of the RfaH-containing coupled TTCs in the loading state obtained using nucleic-acid scaffolds having mRNA spacers of 7, 8, 9, and 10 codons is identical (Figs. S2f); the differences in mRNA spacer length are accommodated through compaction and disorder of mRNA within the TEC RNA-exit channel (Fig. S6a), as observed previously for structures of NusG-containing coupled TTCs^32^. The spatial relationship of the TEC relative to the ribosome is similar, but not identical, to that in structures of NusG-coupled TTCs (Fig. S7a,c-d)^31-32^. The orientation is most similar to that in structures of NusG-coupled TTCs referred to as "subclass B1," representing one of the three--the least populated of the three--main subclasses of NusG-coupled TTCs (Fig. S7a,c-d)^32^.

**Fig. 3.**
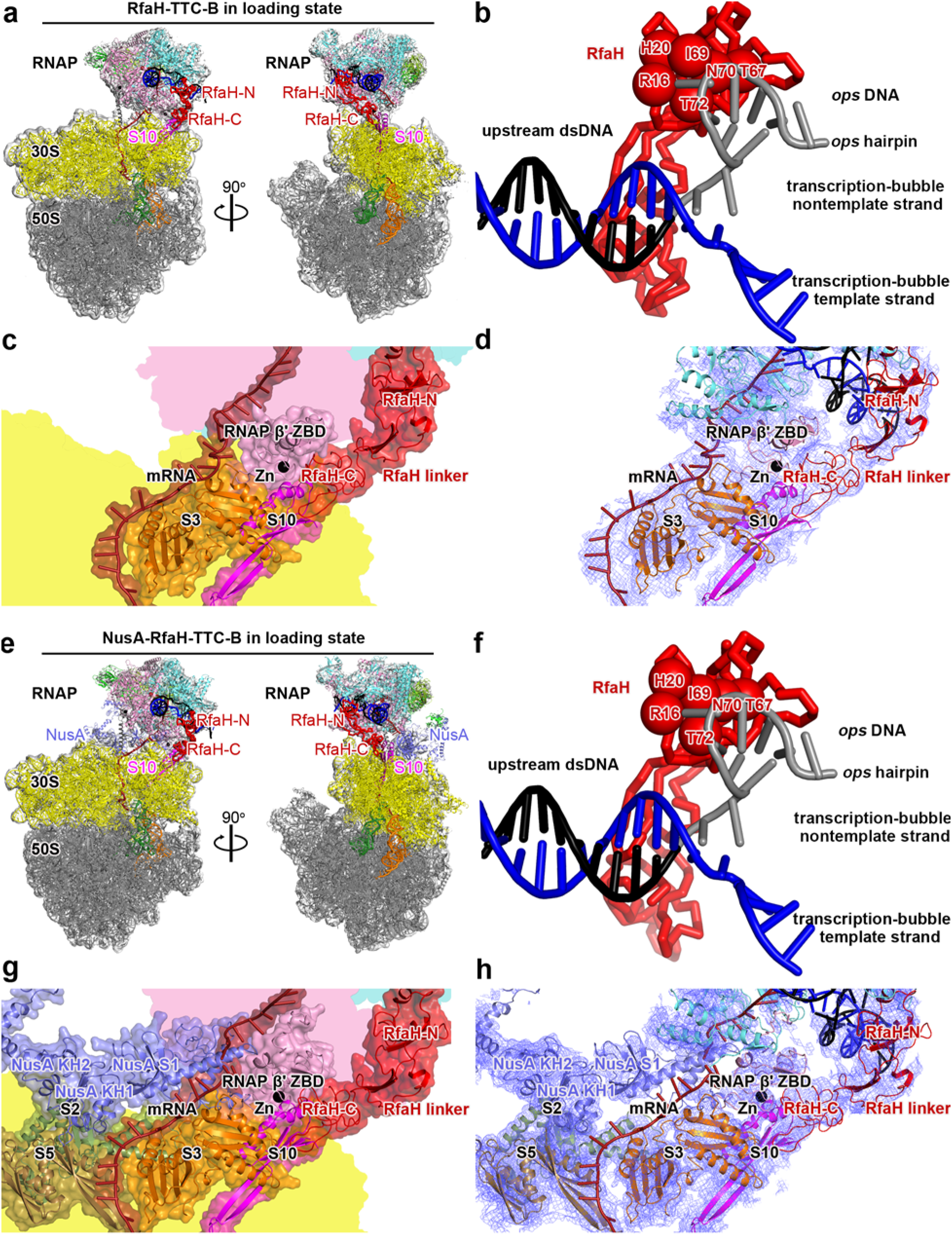
Structures of RfaH-containing coupled TTCs in the loading state. **(a)** Structure of RfaH-TTC-B^loading^ (n = 8; Table S1). Views and colors as in Fig. 2a. **(b)** Protein-DNA interactions between RfaH-N and *ops-*site DNA in RfaH-TTC-B^loading^. **(c)** RNAP-ribosome interface and RfaH bridging in RfaH-TTC-B^loading^ (n = 8; identical interfaces for n = 7, 8, 9, and 10). RNAP β’ zinc binding domain (ZBD, pink; Zn^2+^ ion as black sphere) interacts with ribosomal protein S3 (orange) and mRNA (brick red). RfaH (red) bridges RNAP and ribosome, with RfaH-N interacting with RNAP and RfaH-C interacting with ribosomal protein S10 (magenta). Portions of RNAP β’, β, and ribosome 30S not involved in interactions are shaded pink, cyan, and yellow, respectively. **(d)** As c, showing cryo-EM density as blue mesh. **(e)** Structure of NusA-RfaH-TTC-B^loading^ (n = 8; Table S1). NusA in light blue. Other colors as in a. **(f)** Protein-DNA interactions between RfaH-N and *ops-*site DNA in NusA-RfaH-TTC-B^loading^. **(g)** RNAP-ribosome interface, RfaH bridging, and NusA binding in NusA-RfaH-TTC-B^loading^ (n = 8; identical interfaces for n = 8, 9, and 10). RfaH (red) bridges RNAP and ribosome, with RfaH-N interacting with RNAP and RfaH-C interacting with ribosomal protein S10 (magenta). NusA (light blue) KH1 domain interacts with ribosomal proteins S5 and S2 (brown and forest green). Portions of RNAP β’, β, and ribosome 30S not involved in interactions are shaded pink, cyan, and yellow, respectively. **(h)** As g, showing cryo-EM density as blue mesh.

In RfaH-TTC-B^loading^, the TEC is oriented such that mRNA proceeds from the TEC RNA-exit channel across the RNA-helicase region^38^ of ribosomal protein S3 in the ribosome 30S head, then enters the ribosome 30S subunit mRNA-entrance channel, and, from there, proceeds to the ribosome active-center P-site (Fig. 3a,c-d). RfaH physically couples the TEC and the ribosome by bridging the TEC and the ribosome, with RfaH-N interacting with RNAP, and with RfaH-C simultaneously interacting with ribosomal protein S10 in the ribosome 30S head (Fig. 3a,c-d). RfaH-N makes protein-protein interactions with the RNAP β’ and β pincer tips and the RNAP lobe that are identical to those in structures of RfaH-TEC complexes in the absence of ribosomes^33^ and makes sequence-specific protein-DNA interactions with *ops*-site DNA--including direct contacts with the 2 bp stem and 2 nt loop of a DNA hairpin present in the nontemplate-strand DNA of the transcription bubble--that are identical to those in structures of RfaH-TEC complexes in the absence of ribosomes^33-34^ (Figs. 3a-b, S1g, S2g). RfaH-C interacts with ribosomal protein S10, making interactions that match those proposed from NMR studies of a complex of RfaH-C and ribosomal protein S10^5^ (Figs. 3a,c-d, S1h-i, S2h-i). The interface between RNAP and ribosome in the RfaH-containing coupled TTCs is small (221 Å^2^ buried surface area) and is limited to interactions between the RNAP β’ ZBD and ribosomal proteins S3 and S10 in the ribosome 30S head, (Fig. 3c-d), similar to interactions in NusG-coupled TTCs^31-32^.

We also obtained structures of RfaH-coupled TTCs in the loading state in the presence of NusA (NusA-RfaH-TTC-B^loading^; Figs. 3e-h; S3; Table S1). Structures of NusA-RfaH-TTC-B^loading^ were obtained using nucleic-acid scaffolds having mRNA spacers of 8, 9, or 10 codons (Fig. S3; Table S1), and the structural organization of complexes obtained using nucleic-acid scaffolds having mRNA spacers of 8, 9, and 10 codons was identical (Fig. S3f), with differences in mRNA spacer length being accommodated through compaction and disorder of mRNA in the TEC RNA-exit channel (Fig. S6b). The spatial relationship of the TEC relative to the ribosome in the presence of NusA (Figs. 4e,g-h) was identical to that in the absence of NusA (Fig. 4a,c-d). The conformation and interactions of RfaH in the presence of NusA (Figs. 4e-h, S3g-i) were identical to those in the absence of NusA (Fig. 4a-d; S1g-i, S2g-i). In structures of NusA-RfaH-TTC-B^loading^, NusA formed a second bridge between the TEC and the ribosome, with the NusA N-terminal domain (NTD) interacting with the RNAP β flap-tip helix (FTH) and αCTD^II^, the NusA AR2 domain interacting with RNAP αCTD^I^, and NusA KH1 domain interacting with ribosomal proteins S2 and S5 in the ribosome 30S body (Fig. 3e,g-h), as observed previously in structures of NusA-, NusG-coupled TTCs^32^ As compared to structure determination of RfaH-TTC-B^loading^, structure determination of NusA-RfaH-TTC-B^loading^ was associated with substantially higher particle populations (54% vs. 20% for n = 8, 45% vs. 15% for n = 9, and 28% vs. 9.3% for n = 10; Table S1),, indicating that NusA functionally stabilizes RfaH-TTC-B^loading^. Similar results indicate that NusA functionally stabilizes NusG-TTC-B^32^.

**Fig. 4.**
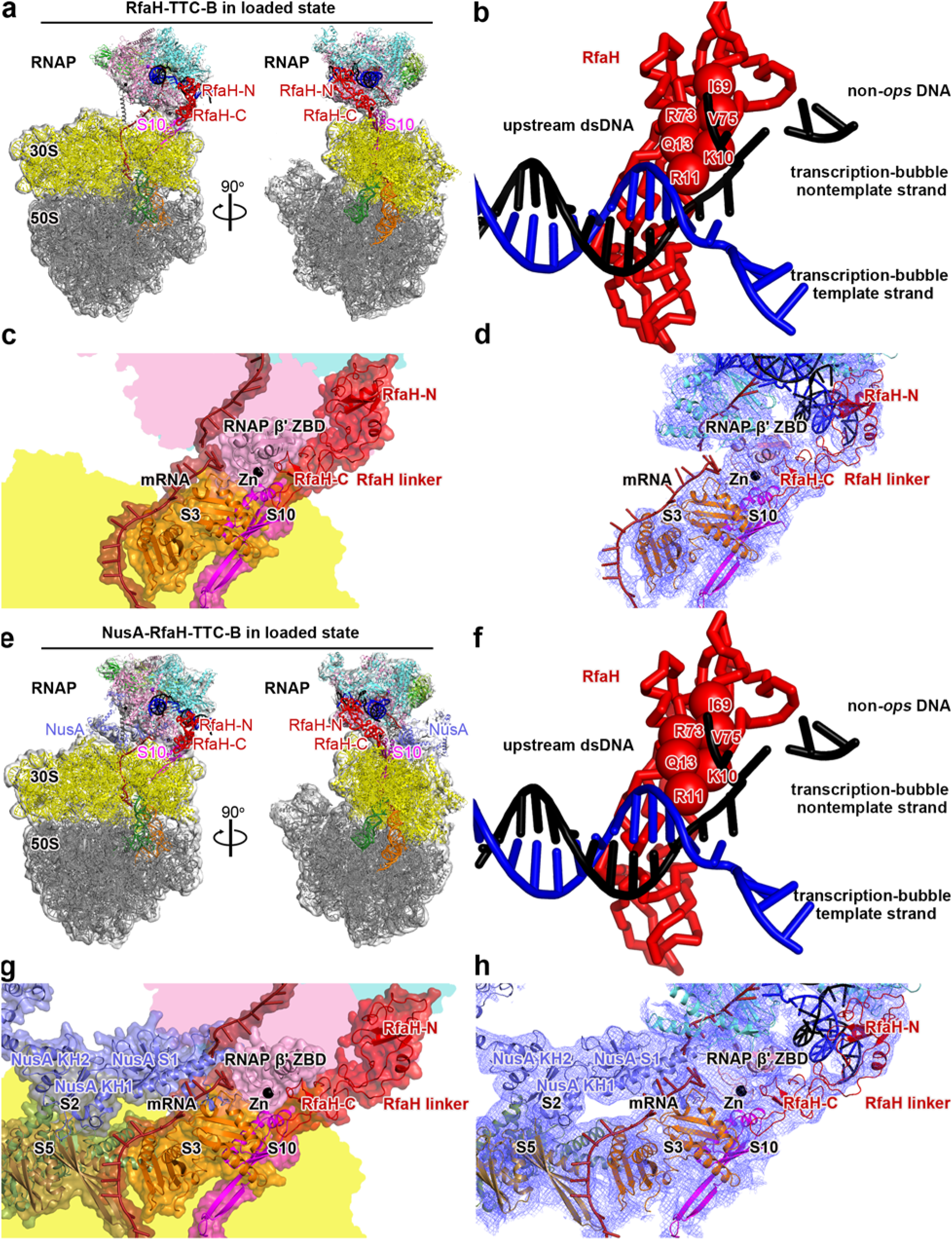
Structures of RfaH-containing coupled TTCs in the loaded state. **(a)** Structure of RfaH-TTC-B^loaded^ (n = 8; Table S1). Views and colors as in Fig. 2a and 3a. **(b)** Interactions between RfaH-N and non-*ops*-site DNA in RfaH-TTC-B^loaded^. **(c)** RNAP-ribosome interface and RfaH bridging in RfaH-TTC-B^loaded^. RNAP β’ zinc binding domain, (ZBD, pink; Zn^2+^ ion as black sphere) interacts with ribosomal protein S3 (orange) and mRNA (brick red). RfaH (red) bridges RNAP and ribosome, with RfaH-N interacting with RNAP and RfaH-C interacting with ribosomal protein S10 (magenta). Portions of RNAP β’, β, and ribosome 30S not involved in interactions are shaded pink, cyan, and yellow, respectively. **(d)** As c, showing cryo-EM density as blue mesh. **(e)** Structure of NusA-RfaH-TTC-B^loaded^ (n = 8; Table S1). NusA in light blue. Other colors as in a. **(f)** Interactions between RfaH-N and non-*ops-*site DNA in NusA-RfaH-TTC-B^loaded^. **(g)** RNAP-ribosome interface, RfaH bridging, and NusA binding in NusA-RfaH-TTC-B^loaded^. RfaH (red) bridges RNAP and ribosome, with RfaH-N interacting with RNAP and RfaH-C interacting with ribosomal protein S10 (magenta). NusA (light blue) KH1 domain interacts with ribosomal proteins S5 and S2 (brown and forest green). Portions of RNAP β’, β, and ribosome 30S not involved in interactions are shaded pink, cyan, and yellow, respectively. **(h)** As g, showing cryo-EM density as blue mesh.

The structures of RfaH-TTC-B^loading^ and NusA-RfaH-TTC-B^loading^, like previously reported structures of NusG-TTC-B and NusA-NusG-TTC-B^32^, appear to be compatible with the ribosome 30S head swivelling and unswivelling^35-37^ that occurs in processive translation. The structures suggest that ribosome 30S head swivelling and unswivelling are accommodated by rotation of the TEC and RfaH with the ribosome 30S head during swiveling and unswivelling, and by flexing and unflexing of the open-rectangular-frame--"pantograph^"32^--structure of NusA during swiveling and unswivelling, as previously suggested for NusG-TTC-B and NusA-NusG-TTC-B^32^. We propose that RfaH-TTC-B^loading^ and NusA-RfaH-TTC-B^loading^, like NusG-TTC-B and NusA-NusG-TTC-B, are functional in processive translation and are complexes that mediate transcription-translation coupling in cells.

Although the structural organization of RfaH-coupled TTCs resembles that of NusG-coupled TTCs^31-32^, there are clear differences in terms of the structural homogeneity of complexes. The range of translational and rotational orientations of the TEC relative to the ribosome to RNAP is much more restricted in RfaH-coupled TTCs than in NusG-coupled TTCs, and the range of conformational states of the RfaH interdomain linker is much more restricted for the RfaH interdomain linker than for the NusG interdomain linker. The difference in conformational homogeneity--with RfaH-coupled TTCs showing much greater structural homogeneity than NusG-coupled TTCs--is manifested in three ways. First, for each mRNA spacer length analyzed, RfaH-coupled TTCs comprise a single structural subclass with a single translational and rotational orientation of the TEC relative to the ribosome (Figs. S2f, S3f, S7a-b), in contrast to NusG-coupled TTCs, which comprise three major subclasses--B1, B2, and B3--and additional minor subclasses (Fig. S7c)^32^. Second, for each mRNA spacer length analyzed, RfaH-coupled TTCs show the same spatial relationship of RfaH-N relative to RfaH-C and the same conformation of the RfaH interdomain linker, in contrast to NusG-coupled TTCs, which show different spatial relationships of NusG-N relative to NusG-C and different conformations of the NusG interdomain linker^32^. Third, for each mRNA spacer length analyzed, cryo-EM density for the RfaH interdomain linker is more ordered than cryo-EM density for the NusG interdomain linker. These differences in conformational homogeneity appear to be attributable to two structural features of RfaH and its protein-protein interactions. First, RfaH-N, but not NusG-N, contains a two-strand β-sheet that departs from the main body of RfaH-N and interacts with the RfaH interdomain linker across its length, stabilizing a single conformational state of the RfaH linker (Figs. 3c-d, 4c-d, S1h-i, S2h-i, S3h-i). Second, RfaH-C, but not NusG-N^32^, makes protein-protein interactions with RNAP β’ ZBD and RNAP β FT that stabilize one spatial relationship of RfaH-C relative to RNAP (Figs, 3c-d, 4c-d).

The higher rigidity of the RfaH interdomain linker, as compared to the NusG interdomain linker, necessarily results in a higher population of states competent with coupling for RfaH than for NusG, and, as such, is expected to result in increased kinetics of coupling during formation of TTCs and decreased kinetics of uncoupling during disassembly of TTCs. We propose that the higher rigidity of the RfaH interdomain linker results in increased kinetics of translation initiation to form TTCs and higher processivity of transcription-coupled translation by the resulting TTCs. Functional and practical implications of this proposal will be discussed below.

### Structures of RfaH-coupled TTCs in the loaded state

Structures of RfaH-coupled TTCs in the loaded state--the state that mediates transcription-translation coupling after the TTC departs from the *ops* site--were obtained using a non-*ops*-site-containing nuclei-acid scaffold, both in the absence and in the presence of NusA (RfaH-TTC-B^loaded^ and NusA-RfaH-TTC-B^loaded^; Figs. 4, S4-S5; Table S1). In RfaH-TTC-B^loaded^ and NusA-RfaH-TTC-B^loaded^, the TEC is positioned relative to ribosome such that RNA proceeds from the TEC RNA-exit channel across the RNA-helicase domain of ribosomal protein S3 in the ribosome 30S subunit, then enters the mRNA entrance channel of the ribosome 30S subunit, and, from there, proceeds to the ribosome active center (Fig. 4a,c-d,e,g-h). RfaH bridges RNAP and the ribosome, and, when present, NusA forms a second bridge between RNAP and the ribosome (Fig. 4 a,c-d,e,g-h). RfaH-N makes the same interactions with the RNAP β’ and β pincer tips and RNAP lobe as in the structures of the corresponding loading complexes, but, due to the absence of an *ops* site, makes none of the sequence-specific protein-DNA interactions by residues 16, 20, 67, 69, 70, and 72 that are made in the loading complexes and appears to make no interactions with the central four nucleotides of the nontemplate strand of the transcription bubble, which are disordered (Fig. 4b,f; compare Fig. 3b,f). The structures of RfaH-TTC-B and NusA-RfaH-TTC-B in the loaded states are superimposable on the corresponding structures in the loading states, with the exception of the interactions of RfaH with DNA (Figs. S4f, S5f). We conclude that there are no substantive differences between RfaH-coupled TTCs in the loading state and in the loaded state, apart from the presence of sequence-specific interactions with DNA in the former and absence in the latter.

## Discussion

Our results show that RfaH couples transcription and translation by physically bridging RNAP and ribosome; show that the range of spatial orientations of RNAP relative to the ribosome in RfaH-coupled TTCs is markedly more restricted than in NusG-coupled TTCs, due to a markedly more rigid interdomain linker in RfaH than in NusG; and show that RfaH-coupled TTCs in the loading and loaded states differ only by the presence of sequence-specific DNA interactions with the *ops* site in the former and the absence in the latter.

Our most notable finding is the markedly greater structural homogeneity and higher conformational rigidity of RfaH-coupled TTCs than of NusG-coupled TTCs. The greater structural homogeneity and higher conformational rigidity are likely to result in increased rate of assembly and decreased rate of disassembly of coupled complexes for RfaH as compared to NusG, and thus are likely to result in more efficient transcription-coupled translation initiation and more processive transcription-coupled translation for RfaH as compared to NusG. The inferred more efficient transcription-coupled translation initiation and more processive transcription-coupled translation for RfaH as compared to NusG likely have both functional consequences and practical consequences.

With respect to functional consequences, we note that the RfaH regulon--the set of transcription units regulated by RfaH--is highly enriched in genes for RNAs that lack Shine-Delgarno sequences^5,26-28,^. We suggest that genes for RNAs that lack Shine-Delgarno sequences are enriched in the RfaH regulon because they require the high efficiency of transcription-coupled translation initiation provided by RfaH. Similarly, we note that the RfaH regulon is highly enriched in transcription units that are exceptionally long (>10,000 bp)^26-28^. We suggest that exceptionally long transcription units are enriched in the RfaH regulon because they require the high processivity of transcription-coupled translation provided by RfaH.

With respect to practical consequences, the greater structural heterogeneity, inferred higher efficiency of transcription-coupled translation initiation, and inferred higher processivity of transcription-coupled translation for RfaH-coupled TTCs, as compared to NusG-coupled TTCs, imply that RfaH-coupled TTCs may be more experimentally tractable for structural and mechanistic studies of transcription-translation coupling. We suggest that structural and mechanistic studies that involve short-lived, low-stability, and/or low-population states--such as cryo-EM structural studies of transcription-coupled translation initiation and transcription-coupled translation, cryo-ET structural studies of *in situ* transcription-coupled translation initiation and transcription-coupled translation, and single-molecule and single-cell functional studies--will be facilitated by targeting RfaH-coupled TTCs, rather than NusG-coupled TTCs.

Our results answer the previously open questions regarding the structural basis of RfaH-dependent transcription-translation coupling, provide a foundation for understanding the role of RfaH in transcription-coupled translation initiation and transcription-coupled translation, and set the stage for, and point to the most effective approaches for, future work to define the mechanisms of transcription-coupled translation initiation and transcription-coupled translation.

## Methods

### *E. coli* RNAP core enzyme, NusA, and 70S ribosomes

*E. coli* RNAP core enzyme, NusA, and 70S ribosomes were prepared as described^32^.

### E. coli RfaH

*E. coli* RfaH^E48A^, an RfaH derivative that contains a substitution in RfaH-N that disrupts interactions between RfaH-N and RfaH-C that inhibit formation of the active conformational state of RfaH-C^5^, was prepared from *E. coli* strain BL21 Star (DE3) (Thermo Fisher) transformed with plasmid pIA763 (encodes *E. coli* RfaH^E48A^ with N-terminal hexahistidine tag^39^), as described^40^.

### *E. coli* tRNA^fMet^

*E. coli* tRNA^fMet^ was purchased (MP Biomedical), dissolved to 100 μM in 5 mM Tris-HCl, pH 7.5, and stored in aliquots at -80°C.

### Nucleic-acid scaffolds

Oligodeoxyribonucleotides and oligoribonucleotides (sequences in Fig. 1b-c) were purchased (Integrated DNA Technologies), PAGE-purified, dissolved in annealing buffer (5 mM Tris-HCl, pH 7.5) to 1 mM, and stored at -80°C in aliquots. Nucleic-acid scaffolds were prepared as described^32^.

### Cryo-EM structure determination: sample preparation

TTCs and EM grids for structure determination were prepared as described^32^, except that NusG was replaced by RfaH^E48A^.

### Cryo-EM structure determination: data collection and data reduction

Cryo-EM data for RfaH-TTC-A (n = 7), RfaH-TTC-B^loading^ (n = 7), RfaH-TTC-B^loading^ (n = 8), RfaH-TTC-B^loading^ (n = 9), RfaH-TTC-B^loading^ (n = 10), NusA-RfaH-TTC-B^loading^ (n = 9), and NusA-RfaH-TTC-B^loaded^ (n = 8) were collected at the Rutgers University Cryo-EM and Nanoimaging Core Facility, using a 200 kV Talos Arctica (FEI/ThermoFisher) electron microscope equipped with a GIF Quantum K2 direct electron detector (Gatan). Data were collected automatically in counting mode, using EPU (FEI/ThermoFisher), a nominal magnification of 130,000x, a calibrated pixel size of 1.038 Å/pixel, and a dose rate of 4.8 electrons/pixel/s. Movies were recorded at 200 ms/frame for 6 s (30 frames), resulting in a total radiation dose of 26.7 electrons/Å^2^. Defocus range was varied between -1.25 µm and -2 µm. For RfaH-TTC-A and RfaH-TTC-B^loading^ (n = 7), a dataset of 1,888 micrographs was recorded from one grid over two days. For RfaH-TTC-B^loading^ (n = 8), RfaH-TTC-B^loading^ (n = 9), RfaH-TTC-B^loading^ (n = 10), NusA-RfaH-TTC-B^loading^ (n = 9), and NusA-RfaH-TTC-B^loaded^ (n = 8), datasets of 9,905, 2,350, 3,055, 1,770, and 3,168 micrographs were recorded from single grids over 2-3 days.

Micrographs were gain-normalized and defect-corrected. Data were processed as summarized in Figs. S1-S3 and S5. Data processing was performed using a Tensor TS4 Linux GPU workstation with four GTX 1080 Ti graphic cards (NVIDIA). Dose weighting motion correction (3×3 tiles; b-factor = 150) were performed using Motioncor2^41^. Contrast-transfer-function (CTF) estimation was performed using CTFFIND-4.1^42^. Subsequent image processing was performed using Relion 3.0^43^. Automatic particle picking with Laplacian-of-Gaussian filtering yielded initial sets of 196,707, 770,444, 228,342, 158,889, 2,034,185, and 259,776 particles for RfaH-TTC-A and RfaH-TTC-B^loading^ (n = 7), RfaH-TTC-B^loading^ (n = 8), RfaH-TTC-B^loading^ (n = 9), RfaH-TTC-B^loading^ (n = 10), NusA-RfaH-TTC-B^loading^ (n = 9), and NusA-RfaH-TTC-B^loaded^ (n = 8), respectively. Particles were extracted into 500×500 pixel boxes and subjected to rounds of reference-free 2D classification and removal of poorly populated classes, yielding selected sets of 10,447, 12,265, 42,674, 7,887, 551,580, and 43,911 particles for RfaH-TTC-A and RfaH-TTC-B^loading^ (n = 7), RfaH-TTC-B^loading^ (n = 8), RfaH-TTC-B^loading^ (n = 9), RfaH-TTC-B^loading^ (n = 10), NusA-RfaH-TTC-B^loading^ (n = 9), and NusA-RfaH-TTC-B^loaded^ (n = 8), respectively. The selected sets were 3D-classified with C1 symmetry, using a *de novo* 3D template created using 3D_initial_model under Relion 3.0. For each complex, classes exhibiting strong, well-defined densities assignable to TEC and having a spatial relationship to density for the ribosome consistent with a TTC were combined and 3D auto-refined using a mask with a diameter of 450 Å. The resulting 3D auto-refined particles were further refined using a soft mask and solvent flattening and were post-processed, yielding reconstructions at 5.5, 5.3, 3.8, 5.7, 6.6, 5.3, and 5.3 Å overall resolution for RfaH-TTC-A (n = 7), RfaH-TTC-B^loading^ (n = 7), RfaH-TTC-B^loading^ (n = 8), RfaH-TTC-B^loading^ (n = 9), RfaH-TTC-B^loading^ (n = 10), NusA-RfaH-TTC-B^loading^ (n = 9), and NusA-RfaH-TTC-B^loaded^ (n = 8), respectively as determined from gold-standard Fourier shell correlation (FSC; Figs. S1-S3, S5; Table S1). The initial atomic models for TTCs were built as described^32^, except that RfaH segments from a cryo-EM structure of *E. coli* RfaH bound to an *E. coli* TEC bound (PDB 6C6S)^44^ were used for docking in place of NusG segments. Refinement of the initial models was performed as described^32^. The initial models were subjected to rounds of manual refinement using Coot^45^ and auto-refinement using Phenix real-space refinement^46^.

Cryo-EM data for NusA-RfaH-TTC-B^loading^ (n = 8) were collected the National Center for CryoEM Access and Training, using a 300 kV Titan Krios (FEI/ThermoFisher) electron microscope equipped with a Gatan K2 Summit direct electron detector (Gatan). Data were collected automatically in counting mode, using Leginon^47^, a nominal magnification of 81,000x and a calibrated pixel size of 1.069 Å/pixel. Movies were recorded at 30 ms/frame for 3 s (100 frames), resulting in a total radiation dose of 45 electrons/Å^2^. Defocus range was varied between -0.8 µm and -2.5 µm. A dataset of 13,660 micrographs was recorded from one grid over 2 days. Data were processed as above, yielding a final set of 2,415,290 particles that produced a reconstruction of 3.4 Å overall resolution (Fig. S3).

Cryo-EM data for NusA-RfaH -TTC-B^loading^ (n = 10) were collected the Stanford-SLAC Cryo-EM Center, using a 300 kV Titan Krios (FEI/ThermoFisher) electron microscope equipped with a Gatan K3 Quantum direct electron detector (Gatan). Data were collected automatically in counting mode using EPU (FEI/ThermoFisher), a nominal magnification of 81,000x and a calibrated pixel size of 1.069 Å/pixel. Movies were recorded at 30 ms/frame for 3 s (100 frames), resulting in a total radiation dose of 45 electrons/Å^2^. Defocus range was varied between -0.8 µm and -2.5 µm. A dataset of 12,096 micrographs were recorded from one grid over 2 days. Data were processed as above, yielding a final set of 48,359 particles that produced a reconstruction of 3.1 Å overall resolution (Fig. S3).

Cryo-EM data for RfaH-TTC-B^loaded^ (n = 8) were collected at the Pacific Northwester Cryo-EM Center, using a 300 kV Titan Krios (FEI/ThermoFisher) electron microscope equipped with a Gatan K3 direct electron detector (Gatan). Data were collected automatically in counting mode, using EPU (FEI/ThermoFisher), a nominal magnification of 81,000x, a calibrated pixel size of 1.069 Å/pixel, a dose rate of 30 electrons/pixel/s, and 30 eV energy slit. Movies were recorded at 50 ms/frame for 2.5 s (50 frames), resulting in a total radiation dose of 65 electrons/Å^2^. Defocus range was varied between -0.8 µm and -2.5 µm. A dataset of 7,071 micrographs was recorded from one grid over 2 days. Data were processed as above, yielding a final set of 1,081,695 particles that produced a reconstruction of 3.2 Å overall resolution (Fig. S4).

Structure visualization was performed using Chimera^46^, Coot^45^, and PyMOL (Schrödinger). Final atomic density maps and atomic coordinates for RfaH-TTC-A (n = 7), RfaH-TTC-B^loading^ (n = 7), RfaH-TTC-B^loading^ (n = 8), RfaH-TTC-B^loading^ (n = 9), RfaH-TTC-B^loading^ (n = 10), NusA-RfaH-TTC-B^loading^ (n = 8), NusA-RfaH-TTC-B^loading^ (n = 9), NusA-RfaH-TTC-B^loading^ (n = 10), RfaH-TTC-B^loaded^ (n = 8), and NusA-RfaH-TTC-B^loading^ (n = 8) were deposited in the Protein Data Bank and the Electron Microscopy Data Bank with accession codes 8UPO and EMD-42453, 8UPR and EMD-42454, 8UQP and EMD-42477, 8URH and EMD-42492, 8URX and EMD-42503, 8UR0 and EMD-42479, 8URI and EMD-42493, 8URY and EMD-42504, 8UQL and EMD-42473, and 8UQM and EMD-42474, respectively..

## Acknowledgments

We thank the Rutgers CryoEM and Nanoimaging Facility, the National Center for CryoEM Access and Training (supported by NIH grant GM129539, Simons Foundation grant SF349247, and New York state grants), the Stanford-SLAC Cryo-EM Center (supported by NIH grant GM129541), and the Pacific Northwest Cryo-EM Center (supported by NIH grant GM129547 and Department of Energy Environmental Molecular Sciences Laboratory) for microscope access. We thank Emre Firlar for technical assistance with cryo-EM data collection.

## Author contributions

V.M. and R.H.E. designed experiments. V.M. and G.B. prepared proteins and nucleic acids. V.M, C.W., and J.T.K. performed cryo-EM data collection. V.M., C.W., and R.H.E. analyzed data. C.W. and R.H.E. prepared figures. R.H.E. wrote the manuscript.

## Data availability statement

Cryo-EM maps have been deposited in the Electron Microscopy Database (EMDB accession codes EMD-42453, EMD-42454, EMD-42473, EMD-42474, EMD-42477, EMD-42479, EMD-42492, EMD-42493, EMD-42503, and EMD-42504), and atomic coordinates have been deposited in the Protein Database (PDB accession codes 8UPO, 8UPR, 8UQL, 8UQM, 8UQP, 8UR0, 8URH, 8URI, 8URX, and 8URY). Unique biological materials will be made available to qualified investigators on request.

## Supplementary Figure Legends

**Fig. S1.**
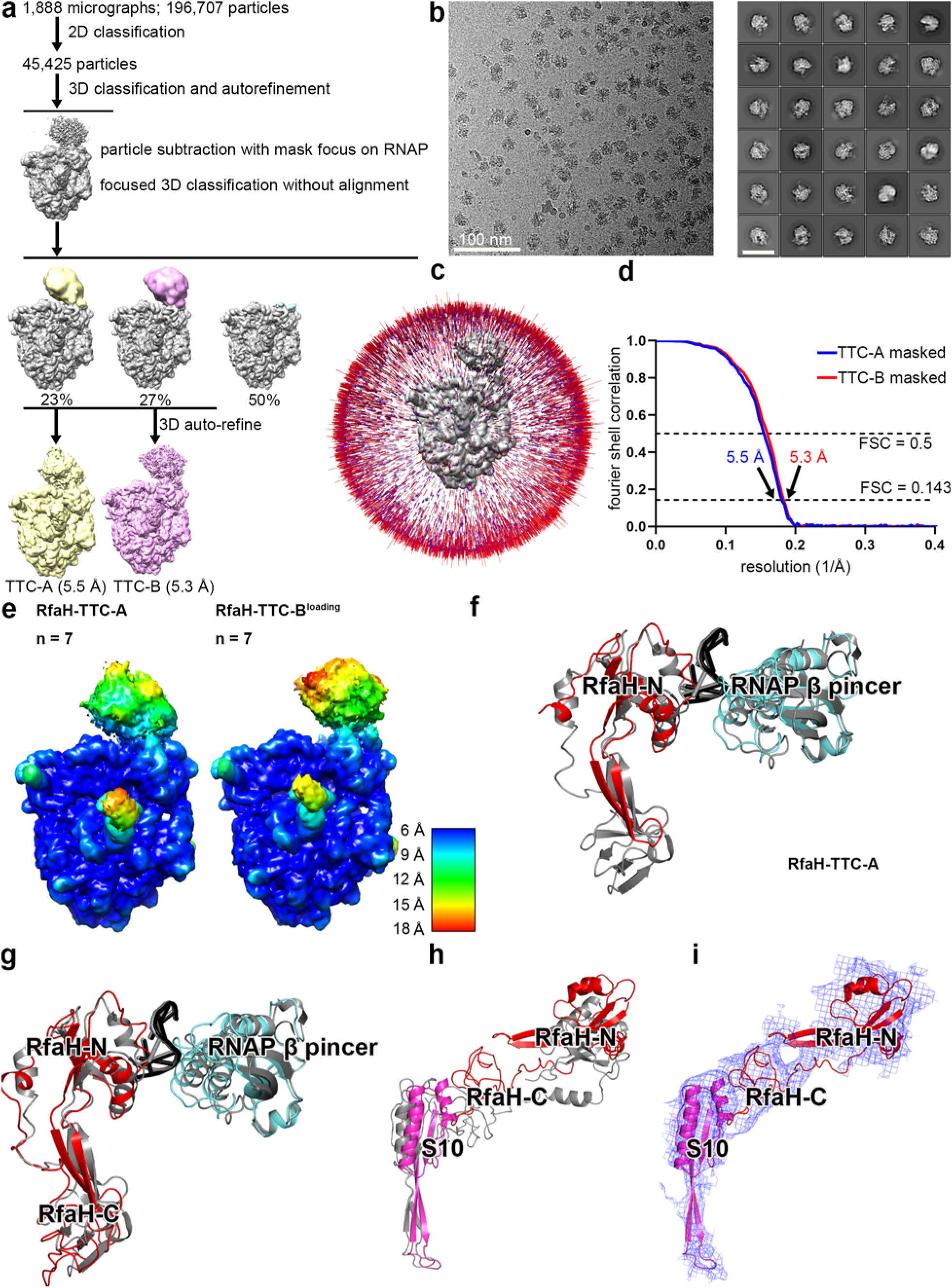
Structure determination: RfaH-TTC-A (n = 7) and RfaH-TTC-B^loading^ (n = 7) **(a)** Data processing scheme (Table S1). **(b)** Representative electron micrograph and 2D class averages (50 nm scale bar in right subpanel). **(c)** Orientation distribution. **(d)** Fourier-shell-correlation (FSC) plot. **(e)** EM density maps colored by local resolution. View orientation as in Figs. 2a and 3a, left. **(f)** Superimposition of RfaH (red) RNAP β’ pincer tip (cyan), and *ops*-site nontemplate-strand DNA in RfaH-TTC-A (n = 7) on corresponding components of RfaH-TEC (PDB 6C6S^33^; gray).. **(g)** Superimposition of RfaH (red), RNAP β’ pincer tip (cyan), and *ops*-site nontemplate-strand DNA in RfaH-TTC-B^loading^ (n = 7) on corresponding components of RfaH-TEC (PDB 6C6S^33^; gray). **(h)** Superimposition of RfaH (red) and ribosomal protein S10 (magenta) in RfaH-TTC-B^loading^ (n = 7) on NusG and ribosomal protein S10 of NusA-NusG-TTC-B subclass B1 (PDB 6X6T^32^; gray. **(i)** EM density (blue mesh) and fit for RfaH (red ribbons) and ribosomal protein S10 (magenta ribbons) in RfaH-TTC-B^loading^ (n = 7).

**Fig. S2.**
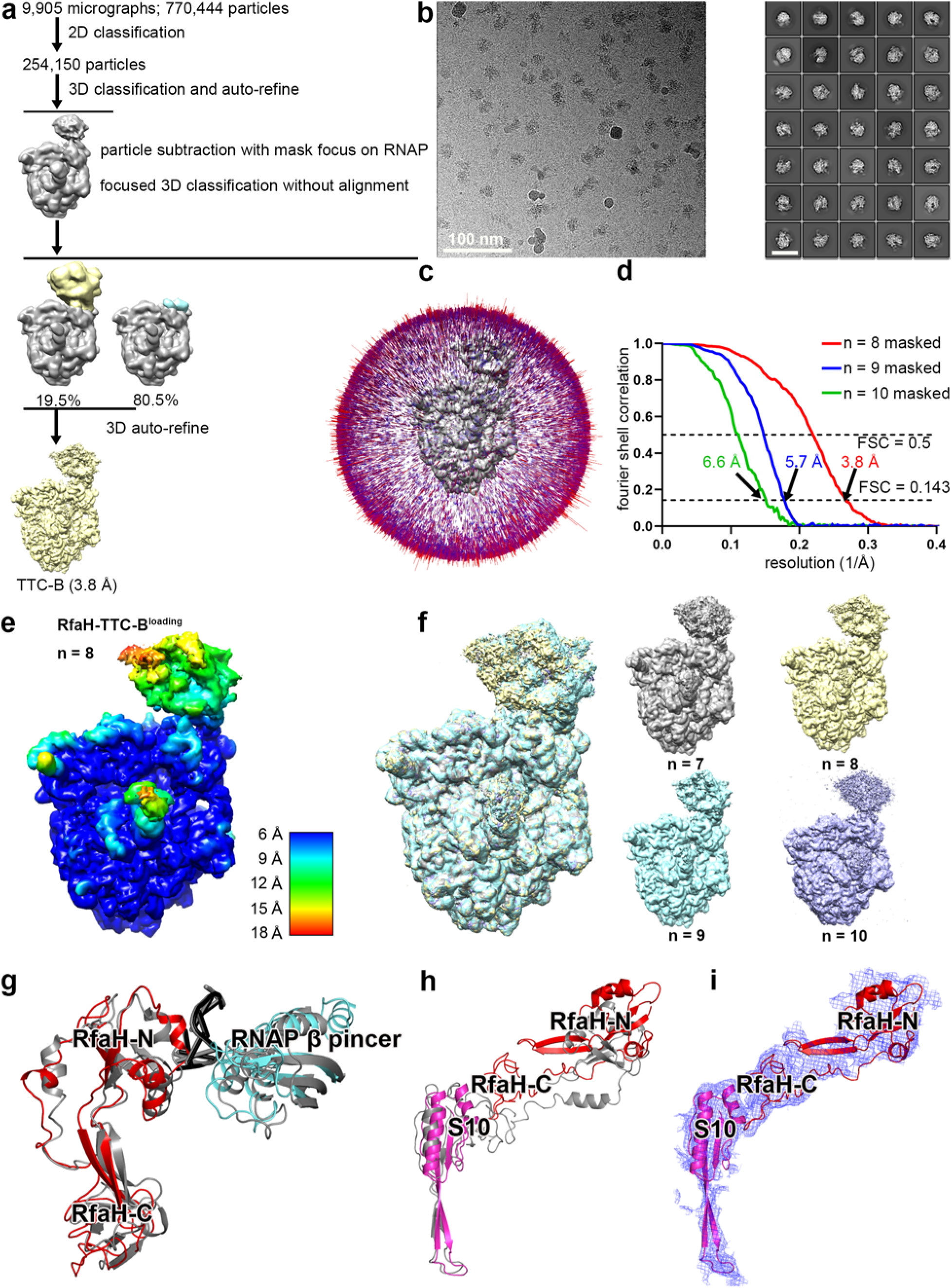
Structure determination: RfaH-TTC-B^loading^ (n = 8, 9, and 10) **(a)** Data processing scheme (Table S1). **(b)** Representative electron micrograph and 2D class averages (50 nm scale bar in right subpanel). **(c)** Orientation distribution. **(d)** Fourier-shell-correlation (FSC) plot. **(e)** EM density maps colored by local resolution. View orientation as in Fig. 3a, left. **(f)** EM density maps of RfaH-TTC-N^loading^ obtained using nucleic-acid scaffolds with n = 7, 8, 9, or 10 (superimposition at left; individual EM maps and color scheme at right). **(g)** Superimposition of RfaH (red), RNAP β’ pincer tip (cyan), and *ops*-site nontemplate-strand DNA in RfaH-TTC-B^loading^ (n = 8) on corresponding components of RfaH-TEC (PDB 6C6S^33^; gray). **(h)** Superimposition of RfaH (red) and ribosomal protein S10 (magenta) in RfaH-TTC-B^loading^ (n = 8) on NusG and ribosomal protein S10 of NusA-NusG-TTC-B subclass B1 (PDB 6X6T^32^; gray. **(i)** EM density (blue mesh) and fit for RfaH (red ribbons) and ribosomal protein S10 (magenta ribbons) in RfaH-TTC-B^loading^ (n = 8).

**Fig. S3.**
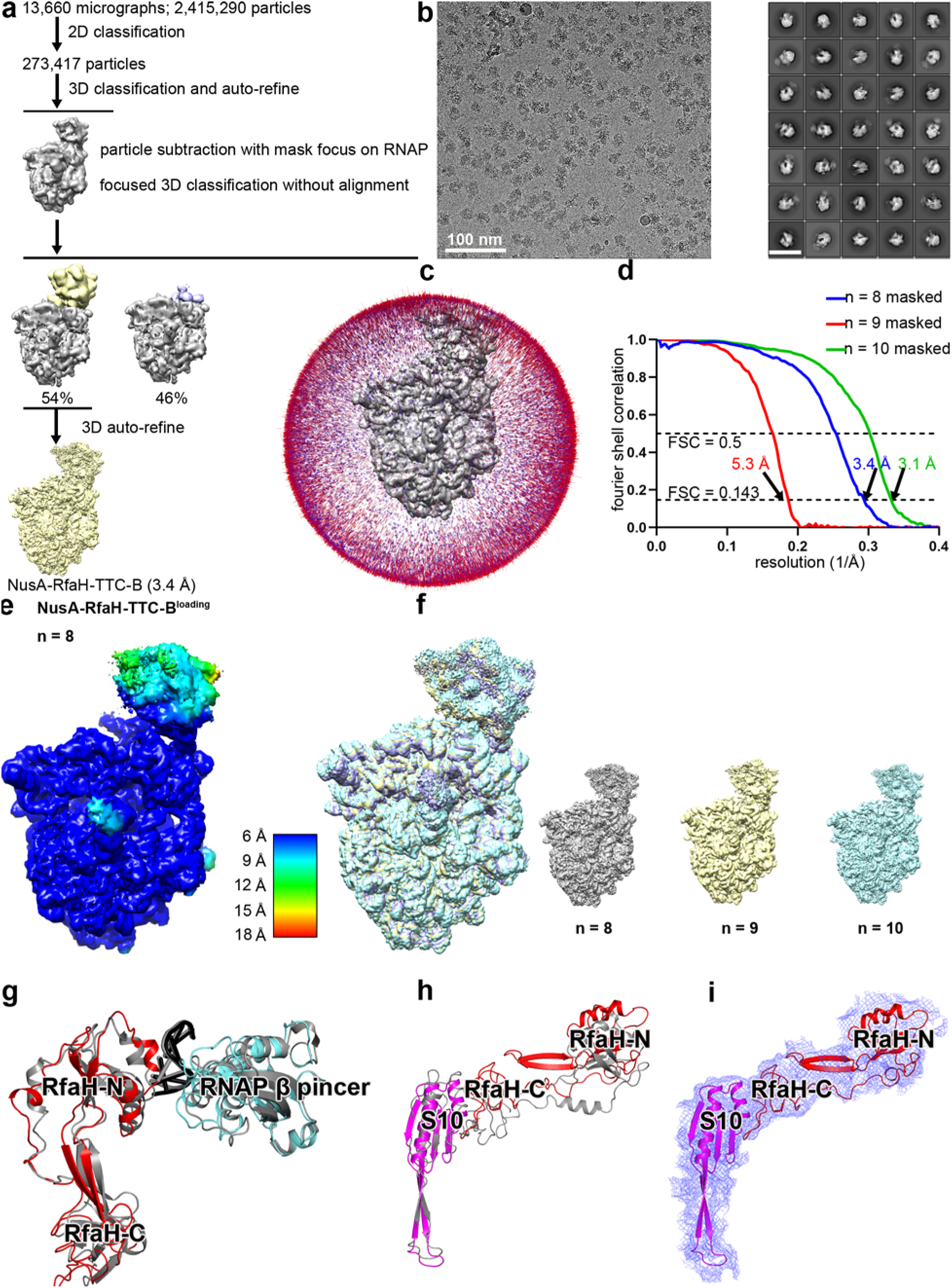
Structure determination: NusA-RfaH-TTC-B^loading^ (n = 8, 9, and 10) **(a)** Data processing scheme (Table S1). **(b)** Representative electron micrograph and 2D class averages (50 nm scale bar in right subpanel). **(c)** Orientation distribution. **(d)** Fourier-shell-correlation (FSC) plot. **(e)** EM density maps colored by local resolution. View orientation as in Fig. 4e, left. **(f)** EM density maps of NusG-RfaH-TTC-N^loading^ obtained using nucleic-acid scaffolds with n = 8, 9, or 10 (superimposition at left; individual EM maps and color scheme at right). **(g)** Superimposition of RfaH (red), RNAP β’ pincer tip (cyan), and *ops*-site nontemplate-strand DNA in NusA-RfaH-TTC-B^loading^ (n = 8) on corresponding components of RfaH-TEC (PDB 6C6S^33^; gray). **(h)** Superimposition of RfaH (red) and ribosomal protein S10 (magenta) in NusA-RfaH-TTC-B^loading^ (n = 8) on NusG and ribosomal protein S10 of NusA-NusG-TTC-B subclass B1 (PDB 6X6T^32^; gray). **(i)** EM density (blue mesh) and fit for RfaH (red ribbons) and ribosomal protein S10 (magenta ribbons) in NusA-RfaH-TTC-B^loading^ (n = 8).

**Fig. S4.**
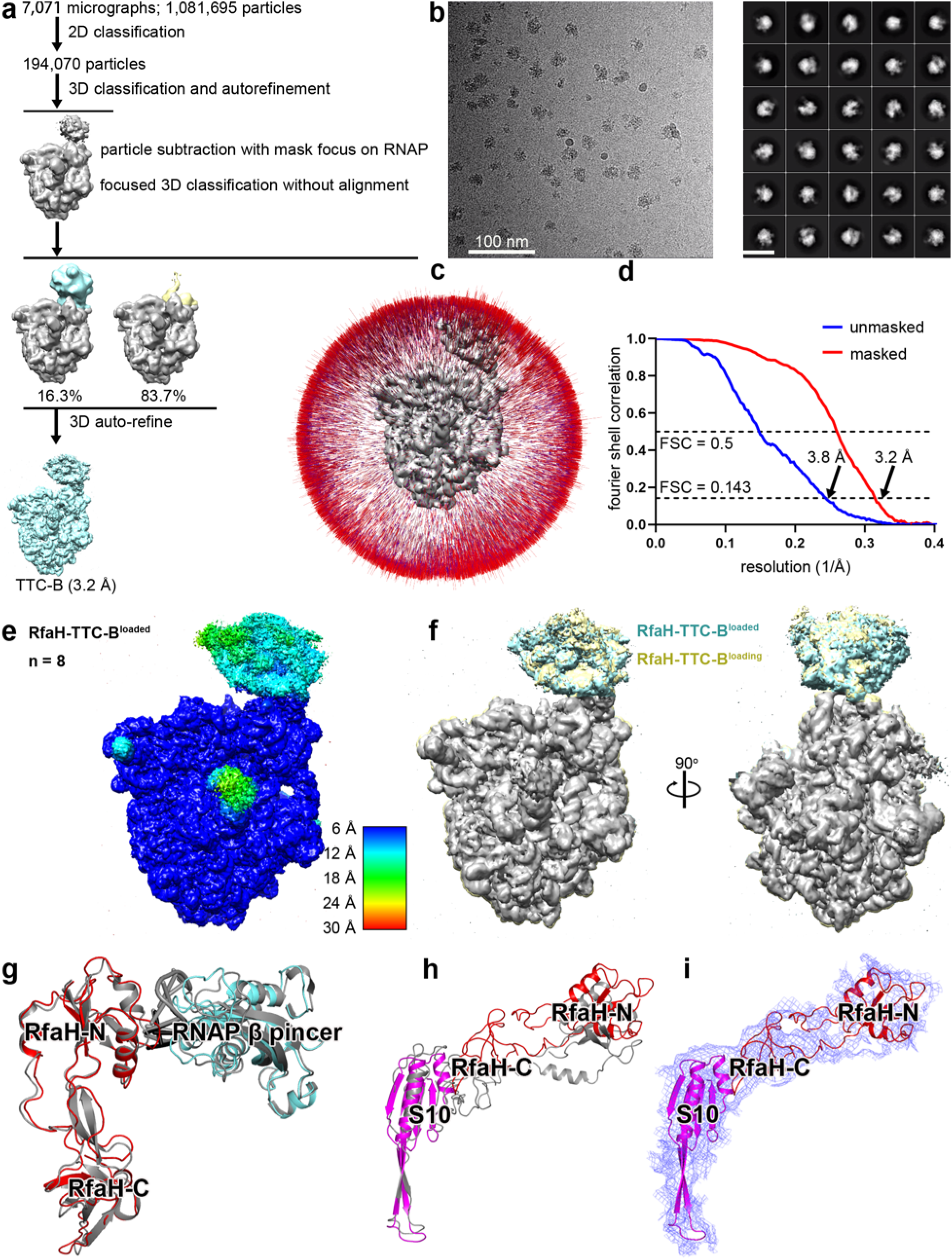
Structure determination: RfaH-TTC-B^loaded^ (n = 8) **(a)** Data processing scheme (Table S1). **(b)** Representative electron micrograph and 2D class averages (50 nm scale bar in right subpanel). **(c)** Orientation distribution. **(d)** Fourier-shell-correlation (FSC) plot. **(e)** EM density map colored by local resolution. View orientation as in Fig. 4a, left. **(f)** Superimposition of RfaH-TTC-B^loaded^ (n = 8) on RfaH-TTC-B^loading^ (n = 8). **(g)** Superimposition of RfaH (red), RNAP β’ pincer tip (cyan), and *ops*-site nontemplate-strand DNA in RfaH-TTC-B^loaded^ (n = 8) on corresponding components of RfaH-TEC (PDB 6C6S^33^; gray). **(h)** Superimposition of RfaH (red) and ribosomal protein S10 (magenta) in RfaH-TTC-B^loaded^ (n = 8) on NusG and ribosomal protein S10 of NusA-NusG-TTC-B subclass B1 (PDB 6X6T^32^; gray.**(i)** EM density (blue mesh) and fit for RfaH (red ribbons) and ribosomal protein S10 (magenta ribbons) in RfaH-TTC-B^loaded^ (n = 8).

**Fig. S5.**
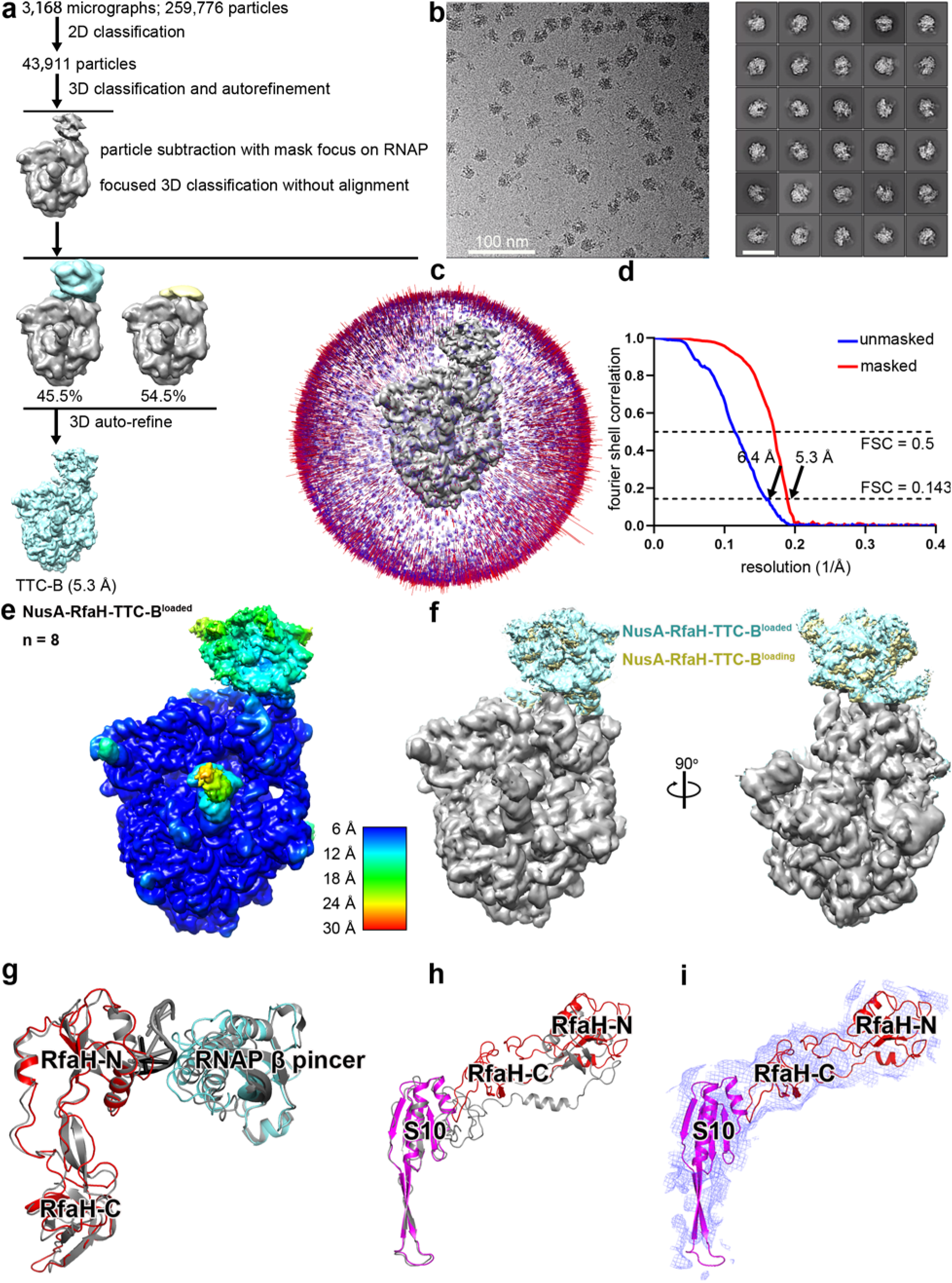
Structure determination: NusA-RfaH-TTC-B^loaded^ (n = 8) **(a)** Data processing scheme (Table S1). **(b)** Representative electron micrograph and 2D class averages (50 nm scale bar in right subpanel). **(c)** Orientation distribution. **(d)** Fourier-shell-correlation (FSC) plot. **(e)** EM density map colored by local resolution. View orientation as in Fig. 4a, left. **(f)** Superimposition of NusA-RfaH-TTC-B^loaded^ (n = 8) on NusA-RfaH-TTC-B^loading^ (n = 8). **(g)** Superimposition of RfaH (red), RNAP β’ pincer tip (cyan), and *ops*-site nontemplate-strand DNA in NusA-RfaH-TTC-B^loaded^ (n = 8) on corresponding components of RfaH-TEC (PDB 6C6S^33^; gray). **(h)** Superimposition of RfaH (red) and ribosomal protein S10 (magenta) in NusA-RfaH-TTC-B^loaded^ (n = 8) on NusG and ribosomal protein S10 of NusA-NusG-TTC-B subclass B1 (PDB 6X6T^32^; gray). **(i)** EM density (blue mesh) and fit for RfaH (red ribbons) and ribosomal protein S10 (magenta ribbons) in NusA-RfaH-TTC-B^loaded^ (n = 8).

**Fig. S6.**
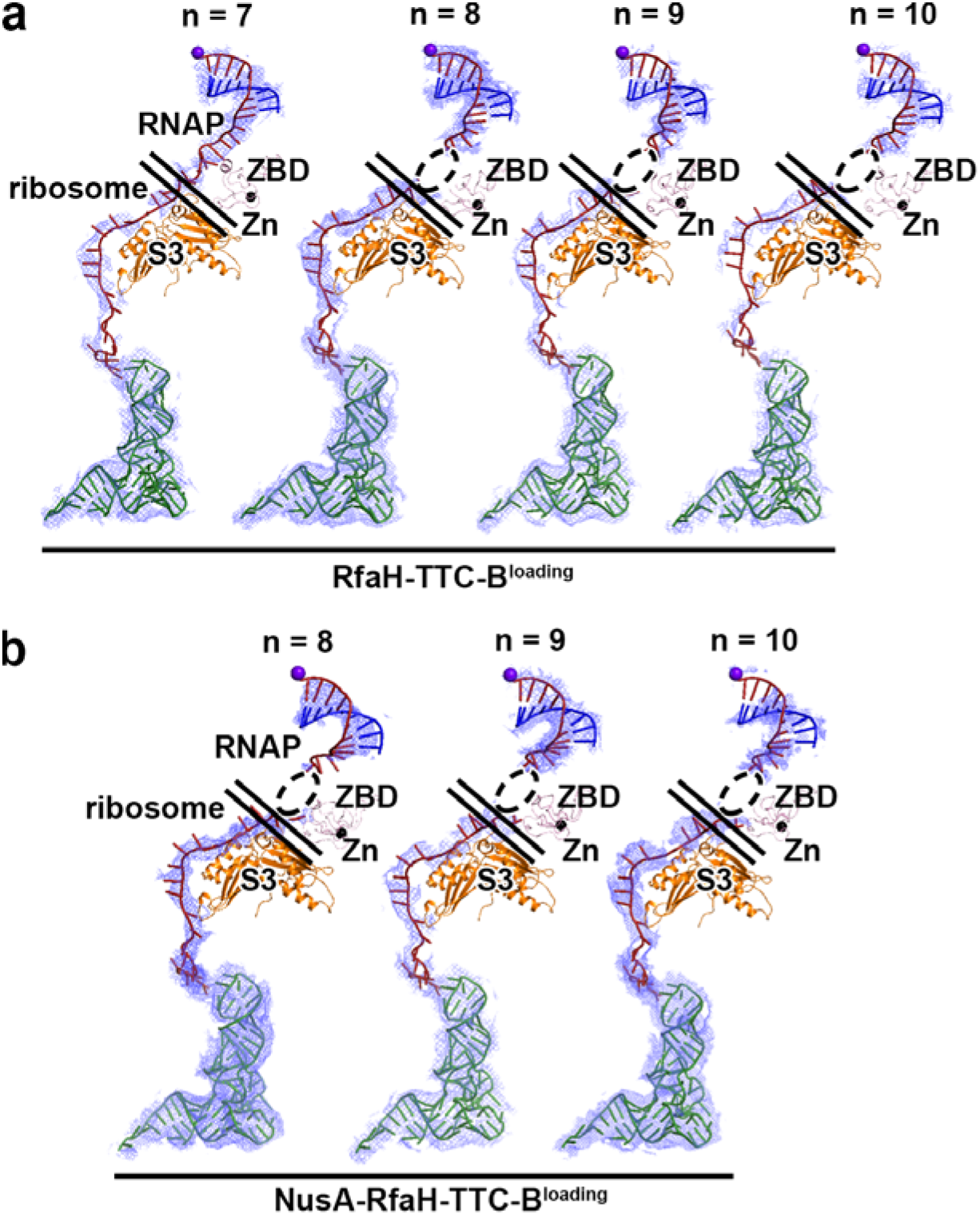
Structures of RfaH-coupled TTCs: accommodation of different mRNA spacer lengths. **(a)** Accommodation of different mRNA spacer lengths of 7, 8, 9, and 10 codons in RfaH-TTC-B (n = 7, 8, 9, and 10). EM density, blue mesh; mRNA, brick-red (disordered mRNA nucleotides indicated by dashed oval); template-strand DNA in RNA-DNA hybrid, blue; RNAP active-center catalytic Mg^2+^, purple sphere; tRNA in ribosome P site, green. Upper and lower black horizontal lines indicate edges of RNAP and ribosome. **(b)** Accommodation of mRNA spacer lengths of 8, 9, and 10 codons in NusA-RfaH-TTC-B (n = 8, 9, and 10). Colors as in a.

**Fig. S7.**
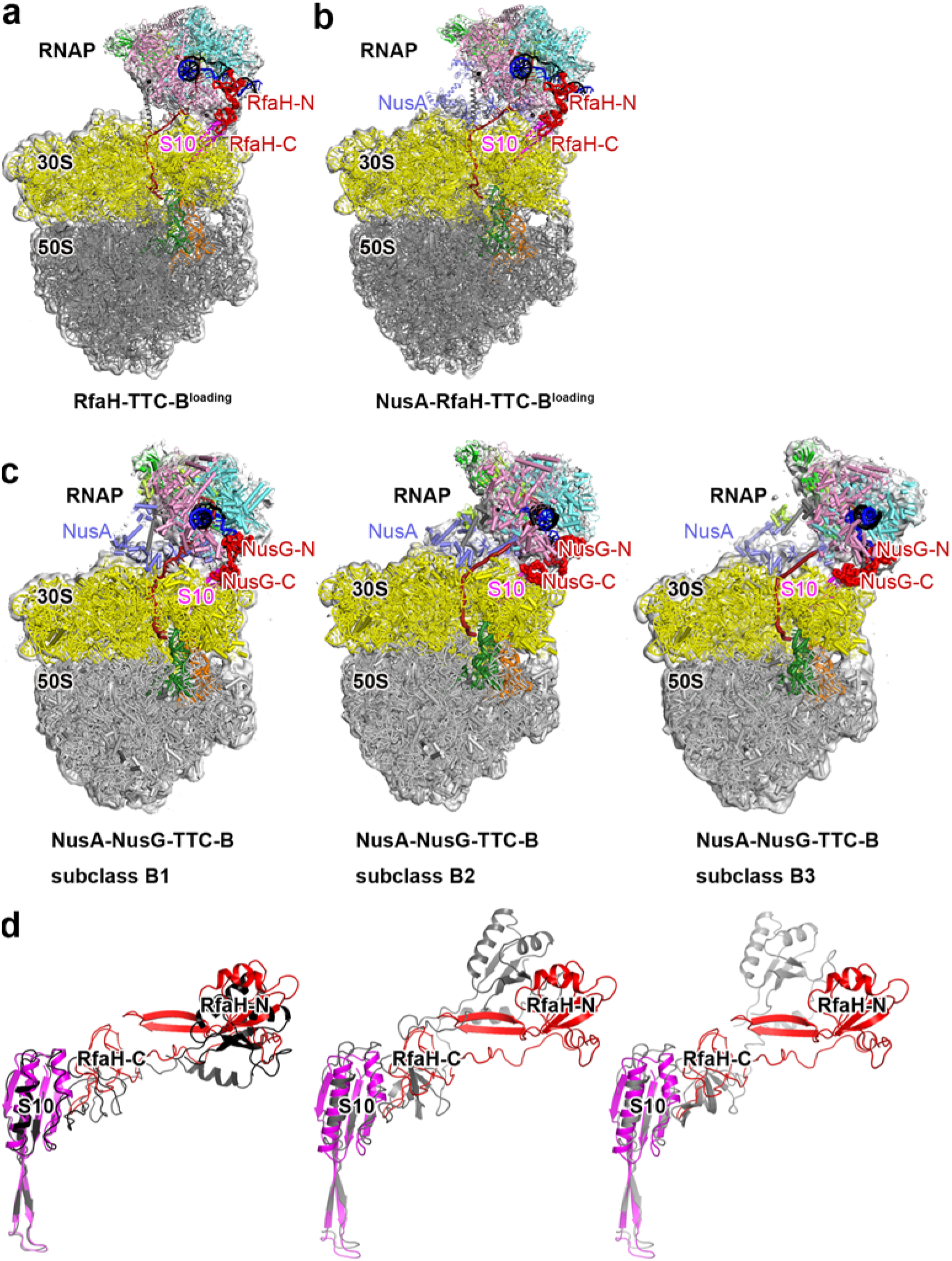
Structures of RfaH-coupled TTCs: comparison of structures of RfaH-coupled TTCs in the loading state to structures of NusG-coupled TTCs. **(a)** Structure of RfaH-TTC-B^loading^ (n = 8) **(b)** Structure of NusA -RfaH-TTC-B^loading^ (n = 8). **(c)** Structure of NusA-NusG-TTC-B subclasses B1, B2, and B3^32.^

**Fig. S8.**
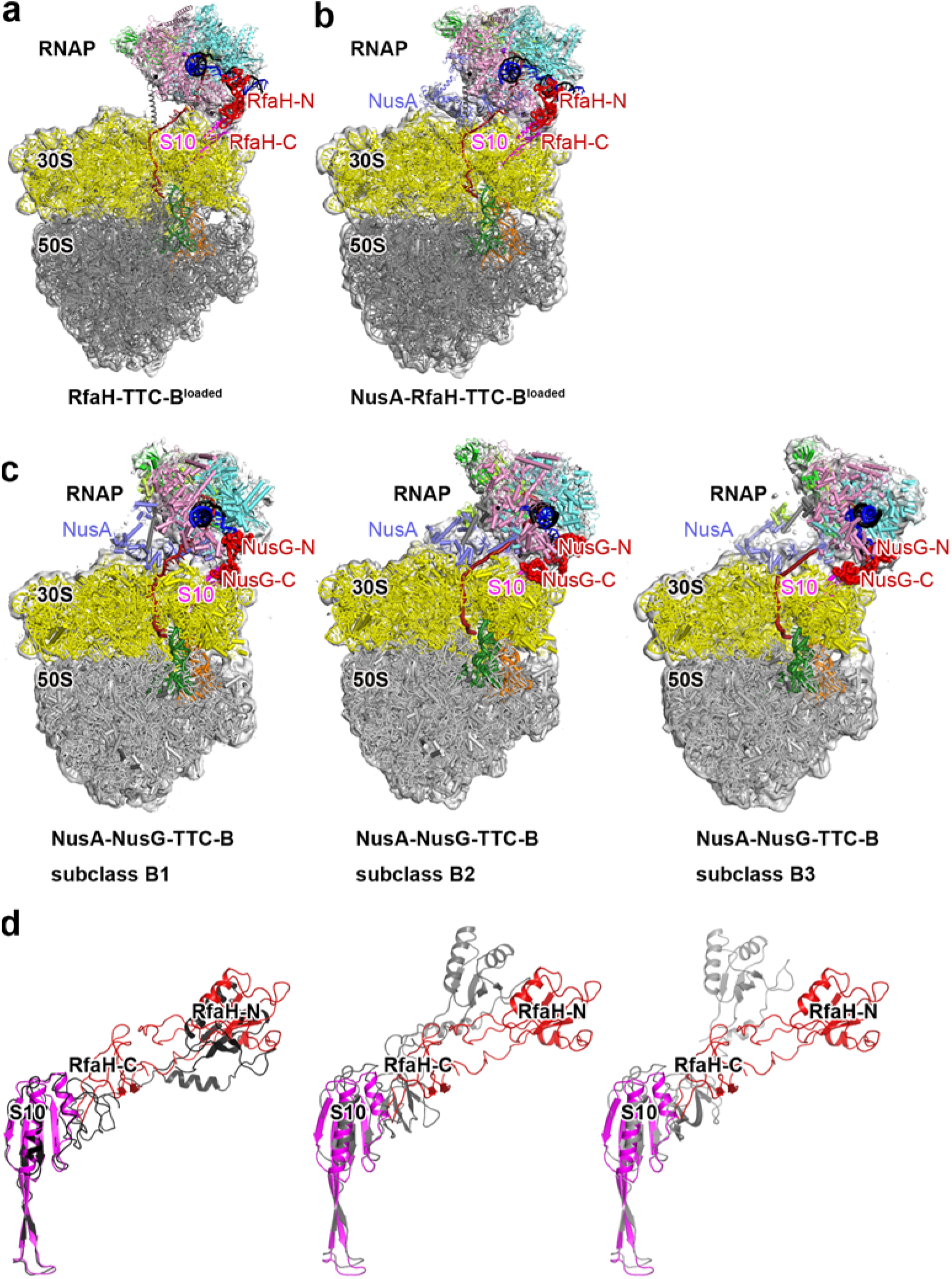
Structures of RfaH-coupled TTCs: comparison of structures of RfaH-coupled TTCs in the loaded state to structures of NusG-coupled TTCs. **(a)** Structure of RfaH-TTC-B^loaded^ (n = 8) **(b)** Structure of NusA -RfaH-TTC-B^loaded^ (n = 8). **(c)** Structure of NusA-NusG-TTC-B subclasses B1, B2, and B3^32.^

**Table S1:** **Cryo-EM structures: RfaH-TTC-A (n = 7), RfaH-TTC-B^loading^ (n = 7, 6, 9, and 10), NusA-RfaH-TTC-B^loading^ (n = 7, 6, 9, and 10), RfaH-TTC-B^loaded^ (n = 8), and NusA-RfaH-TTC-B^loaded^ (n = 8)**

|  | RfaH-TTC-A<br>n = 7<br>(EMDB-42453)<br>(PDB 8UPO) | RfaH-TTC-B<sup>loading</sup><br>n = 7<br>(EMDB-42454)<br>(PDB 8UPR) | RfaH-TTC-B<sup>loading</sup><br>n = 8<br>(EMDB-42477)<br>(PDB 8UQP) | RfaH-TTC-B<sup>loading</sup><br>n = 9<br>(EMDB-42492)<br>(PDB 8URH) |
| --- | --- | --- | --- | --- |
| **Data collection and processing** |  |  |  |  |
| Magnification | 130,000x | 130,000x | 130,000x | 130,000x |
| Voltage (kV) | 200 | 200 | 200 | 200 |
| Electron exposure (e-/Å<sup>2</sup>) | 28 | 28 | 28 | 28 |
| Defocus range (μm) | -1.25 to -2.0 | -1.25 to -2.0 | -1.25 to -2.0 | -1.25 to -2.0 |
| Pixel size (Å) | 1.038 | 1.038 | 1.038 | 1.038 |
| Symmetry imposed | C1 | C1 | C1 | C1 |
| Initial particle images (no.) | 1,888 | 1,888 | 9,905 | 2,350 |
| Final particle images (no.) | 1,720 | 1,720 | 9,873 | 2,212 |
| Map resolution (Å) | 5.5 | 5.3 | 3.8 | 5.7 |
| FSC threshold | 0.143 | 0.143 | 0.143 | 0.143 |
| Map resolution range (Å) | 5.5-15.2 | 5.3-18.6 | 3.8-16.2 | 5.7-16.3 |
| **Refinement** |  |  |  |  |
| Initial models used (PDB codes) | 6C6S, 6VYS | 6C6S, 6VYS | 6C6S, 6VYS | 6C6S, 6VYS |
| Model resolution (Å) | 5.5 | 5.3 | 3.8 | 5.7 |

|  | RfaH-TTC-B<sup>loading</sup><br>n = 10<br>(EMDB-42503)<br>(PDB 8URX) | NusA-RfaH-TTC<br>-B<sup>loading</sup> n = 8<br>(EMDB-42479)<br>(PDB 8UR0) | NusA-RfaH-TTC<br>-B<sup>loading</sup> n = 9<br>(EMDB-42493)<br>(PDB 8URI) | NusA-RfaH-TTC<br>-B<sup>loading</sup> n = 10<br>(EMDB-42504)<br>(PDB 8URY) |
| --- | --- | --- | --- | --- |
| **Data collection and processing** |  |  |  |  |
| Magnification | 130,000x | 81,000x | 130,000x | 81,000x |
| Voltage (kV) | 200 | 300 | 200 | 300 |
| Electron exposure (e-/Å<sup>2</sup>) | 28 | 45 | 28 | 45 |
| Defocus range (μm) | -1.25 to -2.0 | -1.25 to -2.5 | -1.25 to -2.0 | -0.8 to -2.5 |
| Pixel size (Å) | 1.038 | 1.069 | 1.038 | 1.1 |
| Symmetry imposed | C1 | C1 | C1 | C1 |
| Initial particle images (no.) | 3,055 | 13,660 | 1,770 | 12,096 |
| Final particle images (no.) | 2,987 | 13,432 | 1,693 | 11,576 |
| Map resolution (Å) | 6.6 | 3.4 | 5.4 | 3.1 |
| FSC threshold | 0.143 | 0.143 | 0.143 | 0.143 |
| Map resolution range (Å) | 6.6-17.4 | 3.4-14.3 | 5.4-16.5 | 3.1-13.3 |
| **Refinement** |  |  |  |  |
| Initial models used (PDB codes) | 6C6S, 6VYS | 6C6S, 6VYS,<br>6X6T | 6C6S, 6VYS,<br>6X6T | 6C6S, 6VYS,<br>6X6T |
| Model resolution (Å) | 6.6 | 5.3 | 5.4 | 3.1 |

|  | **RfaH-TTC-B<sup>loaded</sup>** | **NusA-RfaH-TTC** |
| --- | --- | --- |
|  | **n = 8** | **-B<sup>loaded</sup> n = 8** |
|  | **(EMDB-42473)** | **(EMDB-42474)** |
|  | **(PDB 8UQL)** | **(PDB 8UQM)** |
| **Data collection and processing** |  |  |
| Magnification | 81,000x | 130,000x |
| Voltage (kV) | 300 | 200 |
| Electron exposure (e-/Å<sup>2</sup>) | 65 | 28 |
| Defocus range (µm) | -0.8 to -2.5 | -1.25 to -2.0 |
| Pixel size (Å) | 1.069 | 1.038 |
| Symmetry imposed | C1 | C1 |
| Initial particle images (no.) | 7,071 | 3,168 |
| Final particle images (no.) | 6,986 | 3,023 |
| Map resolution (Å) | 3.2 | 5.3 |
| FSC threshold | 0.143 | 0.143 |
| Map resolution range (Å) | 3.2-14.7 | 5.3-15.2 |
| **Refinement** |  |  |
| Initial models used (PDB codes) | 6C6S, 6VYS | 6C6S, 6VYS, 6X6T |
| Model resolution (Å) | 3.2 | 5.3 |

